# Beyond the gene: contextualizing novel isoforms through pathway analysis in long-read single-cell RNA sequencing

**DOI:** 10.64898/2026.10.14.758627

**Authors:** Patricia Hayes Doyle, Madeline L. Page, Mark T. W. Ebbert

**Author notes:** To whom correspondence should be addressed: Mark T.W. Ebbert.

## Abstract

Differential isoform usage has been reportedly associated with distinct contexts associated with cell-type and disease; however, the mechanistic role that individual isoforms play in these contexts is poorly understood. Long-read single-cell RNA sequencing enables greater resolution into genes and isoforms previously difficult to detect via short-read approaches, including pseudogenes and those with repetitive motifs. This technique, combined with isoform-discovery tools such as Bambu, has yielded thousands of newly discovered isoforms and genes for consideration. The functional relevance of these new isoforms, however, remains enigmatic. Tools to prioritize which of these novel isoforms should be interrogated and to predict their functionality remain elusive. To address this, we developed a computational framework, Centroid-based Isoform Pathway Inference (CIPI), which contextualizes novel genes and isoforms relative to reference genes with similar cell-type expression patterns. Rather than relying solely on host-gene annotation or similarity to known protein domains, CIPI constructs expression-based neighborhoods that, combined with pathway analyses such as GO, KEGG, or Reactome, enable inference of pathway-level contextualization of individual isoforms according to their biologically meaningful neighbors. Combining CIPI with pathway analyses provides a biological context to novel isoforms, enabling a hypothesis-generating approach for prioritizing novel isoforms across the isonome.

## Introduction

In gene-level analyses, pathway enrichment tools including Gene Ontology (GO) and Kyoto Encyclopedia of Genes (KEGG) can help to identify metabolic pathways, subcellular compartments, and biological processes associated with differences between cell types and disease states. These approaches have advanced our understanding of conditions such as neurodegenerative disease, contextualizing lists of differentially expressed genes and implicating immune dysregulation, mitochondrial dysfunction, and changes in metabolic pathways in the central nervous system and in peripheral immune cells^1,2^. GO analyses were recently combined with machine learning capabilities to create DeepGO^3^, which uses an input protein sequence to assign functional annotations based on homology to known, experimentally annotated proteins and associated GO terminology. While a tool of this design has immense potential for exploring isoform-level differences, its predictions are tied to structural similarity to canonical isoforms alone, limiting its ability to distinguish novel or non-canonical isoform-specific functions. Despite the increasing number of studies highlighting the role of isoform-level diversity, there are no existing pathway analysis tools designed to be used at the isoform level with long-read single-cell data.

One limitation of pathway-based tools is that they were built to be protein-centric and assume a single dominant product per gene. Although gene-level expression and co-expression analyses have been highly informative, they assume that transcript diversity within a gene does not meaningfully alter functional interpretation. This assumption can be inaccurate, given the evidence of alternative isoform expression and divergent isoform function across biology. Approximately 95% of multi-exon genes undergo alternative splicing^4^, producing an average of 14.6 isoforms per gene^5^, though expression abundance appears concentrated among a smaller subset of isoforms^5^. Isoforms within a single gene can encode multiple proteins with drastically different, and even contradictory, functionality within the same pathway, as shown in isoforms of *BCL-X* (*BCL2L1*), which includes a long anti-apoptotic isoform, BCL-X_L_, as well as a short pro-apoptotic isoform, BCL-X_S_^6,7^. These divergent roles cannot be resolved using gene-level pathway enrichment alone.

One of the major challenges in long-read data analysis lies in extracting biological meaning from isoform-level changes in the absence of extensive isoform-level experimental characterization. While isoforms may be differentially expressed or enriched in certain conditions, a crucial juncture is understanding the biological context in which isoform usage is regulated. Interpreting the biological significance of understudied or unknown genes, isoforms, or proteins typically relies on one of three approaches: (1) dissection of known protein domain architecture, (2) use of pathway enrichment analyses, or (3) co-expression networks.

Centroid-based approaches represent the average expression profile of similarly behaving genes within a group (e.g. cell-type, disease, cluster) to act as a representative profile for classification^8,9^, which can be useful to infer functionality. One example in transcriptomics is Weighted Gene Co-expression Network Analysis (WGCNA)^8^, which constructs networks of co-expressed genes and summarizes the module with a dominant expression pattern of the gene set, an eigengene. WGCNA has been used extensively in bulk RNA-seq data, including in Alzheimer’s disease research, where it has identified gene modules tied to neuropathology^10^, cell-type^11–13^, biological processes^2^, and disease status^10^. Recent work has integrated WGCNA with both gene-level and isoform-level single-cell data (hdWGCNA) to extend the module-forming capability^9^, which shows that different isoforms from the same gene can participate in distinctive co-expression modules associated with certain biological trajectories. While these eigenisoforms assemble a module based on networks of co-expression, isoforms are participants within a module, rather than the focal point, making it difficult to infer the functional role of individual isoforms within the module. In addition, these methods do not directly address how co-expression patterns can be used to generate pathway-level inferences for novel isoforms or understudied genes. Thus, the current approaches are limited in their ability to infer novel isoform functionality from single-cell long-read approaches.

To address these limitations, we developed Centroid-based Isoform Pathway Inference (CIPI), an isoform-centered framework that forms reference gene neighborhoods according to similarity in expression patterns to enable pathway analyses. We used isoform-level similarity analyses to show how isoforms within the same gene can have drastically different expression patterns, leading to distinct centroid neighborhoods. We then used these neighborhoods together with pathway enrichment analyses, using known genes to evaluate cell-type-label fit in the context of relevant pathways and to generate pathway enrichment-based hypotheses for novel isoforms based on their proximity to annotated isoforms with similar expression patterns.

## Methods

### Data Preparation

All analyses were performed using the isoform-level single-cell-level expression matrix described in Doyle *et al.*^14^ Corresponding cell-type and sub-cell-type annotations and AutoZI-denoised isoform expression values generated as part of that study were used for downstream analyses. Raw sequencing data were obtained from the NIH Sequence Read Archive (SRA) (accession no. SRR37807425, BioProject no. PRJNA1432415, BioSample no. SAMN56321669).

Data preparation was performed as described in Doyle *et al.*^14^. Briefly, long-read single-cell FASTQ files generated from Dorado underwent quality filtering and pychopper processing. Barcodes underwent targeted correction and rescue using a Levenshtein^15^ distance-based approach, and reads were converted to pseudo-paired-end FASTQ files prior to PIPseeker^16^ processing. Reads were aligned to the GRCh38 reference genome (release 113) using minimap2^17^, and Bambu^18^ (v3.8.1) was used to perform isoform discovery and quantification on the pseudobulk level. Quantification was repeated on the single-cell level, and single-cell matrices were generated. Count matrices were filtered according to the previously described quality metrics: mitochondrial gene percentage <15%, hemoglobin gene percentage <5%, total counts >600 and <11000, unique isoform count >350 and <3000. AutoZI^19^ modeling was then performed as previously described using 20 latent dimensions, a dropout rate of 0.3, 90% of cells used for training data, a learning rate of 0.01, early stopping with patience of 70, and a weight decay of 0.0005.

After modeling, the denoised data were extracted and normalized per 10,000 reads. Latent representations were extracted from the model and used to construct a k-nearest neighbors graph with 20 neighbors per cell, followed by Leiden clustering at resolutions of 0.06 for cell-type and 0.26 for T cell sub-type assignment. Clusters were annotated according to the previously described marker genes for the following cell-types: T cells (*CD3D*, *CD3E*, *CD3G*), CD8 effector T cells (*CD8A*, *CD8B*, *GATA3*, *KLRB1*, *CCL5*), CD4 effector T cells (*CD4*, *IL2RA*, *GATA3*, *AHR*, *TNF*), memory T cells (*CCR7*, *SELL*, *TCF7*), effector-memory transition T cells (*ITGAE*, *LEF1*, *IL2RA*, *CTLA4*, *GATA3*, *IL7R*, *CD27*, *TCF7*), Natural Killer (NK) cells (*GZMB*, *KLRF1*, *NCAM1*, *ITGAM*, *IL2RB*), B cells (*CD22*, *CD79A*, *MS4A1*, *CD19*), monocyte-derived cells (*FCGR2A*, *CLEC7A*, *CD33*, *LILRB4*), and megakaryocytes (*GP1BA*, *MPL*, *ITGA2B*).

### Isoform categorization

Isoforms were split into one of four groups: *reference*, *markerRefIso*, *knownGeneNovelIso*, or *novelGeneNovelIso*. Isoforms were classified as *reference* isoforms if they contained both Ensembl gene (ENSG) and transcript (ENST) identifiers from Ensembl^20^. To establish a positive control dataset for validating our centroid behavior, we utilized the cell-type markers for our major PBMC lineages and T cell subtypes. Our group labeled *markerRefIso* was established as a subset of the *reference* isoforms with those belonging to a pre-defined marker gene set representing major Peripheral Blood Mononuclear Cell (PBMC) lineages and T cell subtypes, as described in Doyle *et al.*^14^. Isoforms categorized as *knownGeneNovelIso* had a known ENSG ID from Ensembl and an isoform ID starting with “BambuTx”, whereas isoforms belonging to the *novelGeneNovelIso* group had a gene ID starting with “BambuGene” and an isoform ID starting with “BambuTx”.

### Expression filtering

To restrict isoforms to those sufficiently supported in our expression matrices, we applied additional filtering criteria to our expression matrix. Isoforms were required to have a non-zero value in ≥10 cells in at least one cell type and to be expressed in ≥1% of cells in at least one cell type. All isoforms from our pre-filtered dataset met these criteria, and all subsequent centroid, similarity, residual, and divergence analyses were performed using this final filtered isoform dataset.

### Cell-type expression summaries

For each isoform, mean expression and isoform detection frequency were independently calculated per cell type. Isoform detection frequency was calculated as the fraction of cells with nonzero expression. To emphasize relative cell-type-specificity rather than absolute expression magnitude, which could be biased by the number of cells detected, each per-cell-type mean expression value was z-score normalized across cell types. Standardization used the cell-type population standard deviation (ddof=0).

### Reference gene centroid construction

*Reference* isoforms were grouped by host gene according to a shared ENSG ID, aggregated by cell type, and their z-scored cell-type expression profiles were averaged to produce a gene-level reference expression profile (hereafter referred to as a “gene centroid”), representing a pseudobulk expression structure for each cell-type per gene. In the resulting matrix, each row corresponded to a gene and each column to a cell type, with values representing the average z-scored expression across all reference isoforms for that gene in that cell type.

### Centroid similarity search and baseline estimation

For isoforms classified as being in the *markerRefIso*, *knownGeneNovelIso*, or *novelGeneNovelIso* group, the similarity of their z-scored cell-type expression profile was compared to those belonging to *reference* gene centroids using a Pearson correlation. For each isoform of interest, the gene centroids with >80% similarity (r>0.8) were ranked by correlation, and up to the top 30 most similar *reference* centroids were assigned as that isoform’s nearest gene centroids, hereby referred to as their “neighbors”. An expected baseline expression profile was computed for each isoform of interest by aggregating the cell-type expression profiles of its nearest centroid neighbors. This baseline profile was computed as a similarity-weighted average of reference centroids, where non-negative correlation coefficients were normalized to add up to one. Isoform-level, cell-type-specific residuals were calculated as the difference between the isoform’s observed z-scored expression and the baseline expectations.

### Positive-control validation

To validate that centroid neighborhoods accurately reflected similar functionality as the isoform of interest, the *markerRefIso* isoforms and their neighbors were used as positive controls. The *reference* gene centroid neighbors for each isoform were ranked in terms of similarity of cell-type enrichment profile as evaluated by Pearson correlation, and the rank of the known host gene was recorded. This was also performed for *knownGeneNovelIso* isoforms.

### Within-host neighbor divergence

For genes with multiple isoforms analyzed, the overlap of gene centroids in CIPI neighborhoods from the same gene were compared using a Jaccard similarity index^21^. All pairwise combinations of isoforms associated with the host gene were evaluated. Lower Jaccard similarity indicated less overlap between reference-gene neighborhoods from isoforms within the same gene.

### Cell-type-specificity

To assess cell-type specificity, we used the τ statistic^22^, Gini coefficient^23,24^, and normalized Shannon entropy^25^ on positive isoform z-scores, to minimize bias obtained from using one measure. τ statistic and Gini coefficient were both used to measure the degree to which expression was concentrated in a limited number of cell types, whereas normalized Shannon entropy provided a measure of expression diversity across cell-types. Gini and Shannon entropy have both demonstrated robustness as a measure of gene expression profiling in previous studies^26,27^. We applied conservative deviation thresholds for decoupling from the host gene’s expression and cell-type specificity, annotating any isoforms that fell above the 90^th^ percentile (q=0.9) for both. Divergent isoform candidates were identified as ones that exhibited a maximum isoform z-score of ≥1.5, high host-gene decoupling (q=0.9), and high τ specificity (q=0.9).

### Goodness-of-fit evaluation

Agreement between an isoform’s expression profile and its centroid-derived baseline was evaluated using Pearson correlation, root mean squared error (RMSE), and the coefficient of determination (R^2^). Pearson correlation quantified the similarity in the pattern of cell-type enrichment. RMSE evaluated how much the observed z-scores deviated from the baseline z-scores. R^2^ quantified the proportion of variance in the observed isoform z-score profile represented by the centroid-derived baseline. Host-isoform decoupling was quantified as one minus the Pearson correlation coefficient (between each isoform’s z-scored cell-type expression profile and its host gene centroid), to reflect how much an isoform’s cell-type expression profile deviated from that of the host gene.

### Functional enrichment

To infer possible biological functions associated with isoform-specific expression patterns, reference gene centroid neighbors for each isoform were mapped to HGNC gene symbols using the Ensembl GRCh38 (release 113)^20^ genome annotation and assembled as a study set input for enrichment analyses for each isoform neighborhood. For all pathway analyses, gene sets were required to map at least three gene symbols prior to undergoing pathway enrichment. Gene Ontology (GO)^28–30^ enrichment analyses were performed using GOATOOLS (v1.1.5)^31^ with the Gene Ontology Basic ontology (go-basic.obo) file and human Gene Ontology annotation file (goa_human.gaf) obtained from the Gene Ontology Consortium (release 2025-10-10)^32^. Enrichment was evaluated across Biological Process (BP), Molecular Function (MF), and Cellular Component (CC) ontologies. To limit cell-type-specific bias and ensure accuracy, enrichment was tested against annotated human genes in the GO database (GOA Human GAF 2.2) represented in the dataset.

Count propagation was enabled, so genes aligned to GO terms were directly annotated as well as their parent terms to emphasize functionality from a broad and specific perspective. Parent terms were also highlighted in plots to show higher-level processes enriched by the isoform neighborhoods. Statistical significance, as calculated using the Benjamini-Hochberg FDR correction^33^, accounted for the likelihood that neighbor genes would be assigned by chance compared to the reference genes in the dataset, with terms retained at an FDR less than 0.05. We then performed a cross-database functional enrichment validation using the g:profiler^34,35^ API (06-15-2025 release) with GO biological pathway (GO:BP), Kyoto Encyclopedia of Genes and Genomes (KEGG)^36^ and Reactome^37–41^ annotations to increase confidence in similarly labeled pathways. g:GOSt performed over-representation analyses using a cumulative hypergeometric test, and statistical significance was assessed with Benjamini-Hochberg FDR multiple testing correction^33^ and a maximum FDR significance threshold at 0.05. As with GOATOOLS, a custom background was used, consisting of the reference centroid genes to prevent FDR inflation by genes not represented in our dataset. To aid in interpretation of GO:BP results, significant terms were annotated with their respective parent-child relationships within the GO hierarchy. A GO term was labeled a “parent” term if at least one of its “child” terms was also significant within the same isoform neighborhood analysis.

## Results

### Overview of CIPI for isoform-level biological context

In this study, we utilized the AutoZI-denoised single-cell expression matrices and cell-type annotations from Doyle *et al.*^14^ To prepare our dataset for pathway enrichment, we averaged expression counts per isoform using all cells within a single cell type and computed the detection frequency of an isoform per cell type. We then performed z-score normalization for isoform cell-type expression profiles to emphasize both cell-type enrichment and relative abundance (**Figure 1a**). Isoforms in our dataset were categorized as being *reference* isoforms, marker isoforms (*markerRefIso)*, new isoforms from new genes (*novelGeneNovelIso)*, or new isoforms from known genes (*knownGeneNovelIso)*. The marker isoforms were from the known gene markers of cell identity used for cell-type assignment by Doyle *et al.*^14^

**Figure 1.**
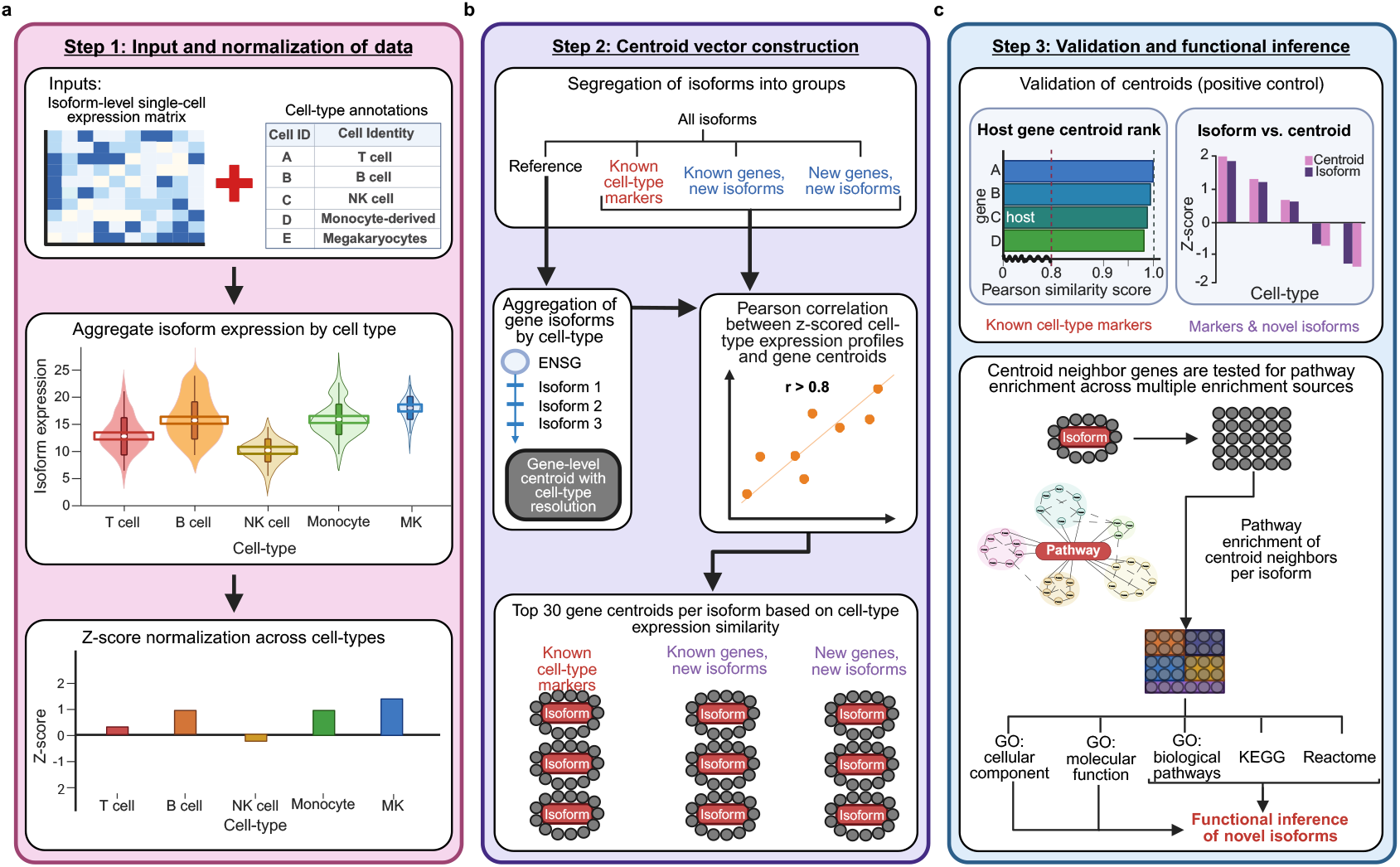
Overview of centroid analysis for cell-type-resolved pathway inference. **(a)** Step 1: A denoised isoform-level single-cell expression object containing layers with cell-type and sub-cell-type annotations was aggregated by cell-type. For each isoform, mean expression and detection frequency were computed according to cell-type expression and means were z-score normalized to emphasize relative cell-type enrichment. **(b)** Step 2: Isoforms were categorized into one of the following groups: *reference*, marker reference (*markerRefIso)*, new isoforms from known genes (*knownGeneNovelIso)* or new isoforms from unknown genes (*novelGeneNovelIso)*. *Reference* isoforms were grouped by host gene (ENSG) to construct gene-level reference centroids, while maintaining z-scores for each cell type. Cell type expression profile similarity from *reference* gene centroids were compared to those from isoforms in the other three groups using Pearson correlation. Gene centroids with high z-score similarity to an isoform (r > 0.8) were ranked according to similarity, and the 30 most similar centroids were assigned as centroid neighbors for each non-reference isoform. **(c)** Step 3: Similarity-weighted baseline expression profiles for each cell-type within a centroid neighborhood were calculated as a prediction of expression per cell-type. Deviations from the centroid-derived baseline were calculated as isoform residuals. Marker reference isoforms (*markerRefIso)* served as positive controls to validate the centroid neighborhood framework and goodness-of-fit between observed isoform expression profiles and predicted centroid baselines. Reference genes within each isoform centroid neighborhood underwent pathway enrichment analyses (Gene Ontology, GO; Kyoto Encyclopedia of Genes and Genomes, KEGG; Reactome) to support pathway inference for unannotated (new) isoforms from new and known genes.

We sought to construct representative centroids that summarized gene-level cell-type tendencies for each gene, to match the expected input for pathway enrichment analyses, without removing cell-type-specific profiles. For each marker isoform and novel isoform, a centroid neighborhood was constructed from the top 30 most similar gene centroids with a similarity of r > 0.8 (**Figure 1b**). These neighborhoods acted as reference units for validation of cell-type assignment and inference of pathways associated with novel and marker isoforms. The marker isoforms were used as positive controls to assess the biological relevance and validity of the centroid neighborhoods. Genes from each neighborhood were used as gene set inputs for pathway enrichment analyses to generate functional hypotheses for novel isoforms (**Figure 1c**).

### Centroid-based references preserve the biological relevance of isoforms not found on the gene level alone

To validate that centroid-based similarity reliably groups genes with similar expression patterns, we used isoforms from recognized immune cell-type marker genes as a positive control. Because our gene centroids represent an average expression profile by cell type across all reference isoforms within a gene, an isoform of interest may not correlate most strongly with its host gene centroid if it exhibits a distinct cell-type-specific expression pattern. Consistent with this, across 173 marker isoforms, the closest centroid was not always that isoform’s host gene. In this set, the host gene had a median rank of 20 in our marker isoform set, being ranked as the top centroid in only 12.7% of isoforms and in the top five centroids for just 28.9% of isoforms (**Figure 2a**). This indicates that individual isoforms exhibit cell-type-expression profiles more similar to those of other genes than the aggregate profile of their host gene, underscoring the limitations of host gene-level analyses for capturing isoform-specific effects.

**Figure 2.**
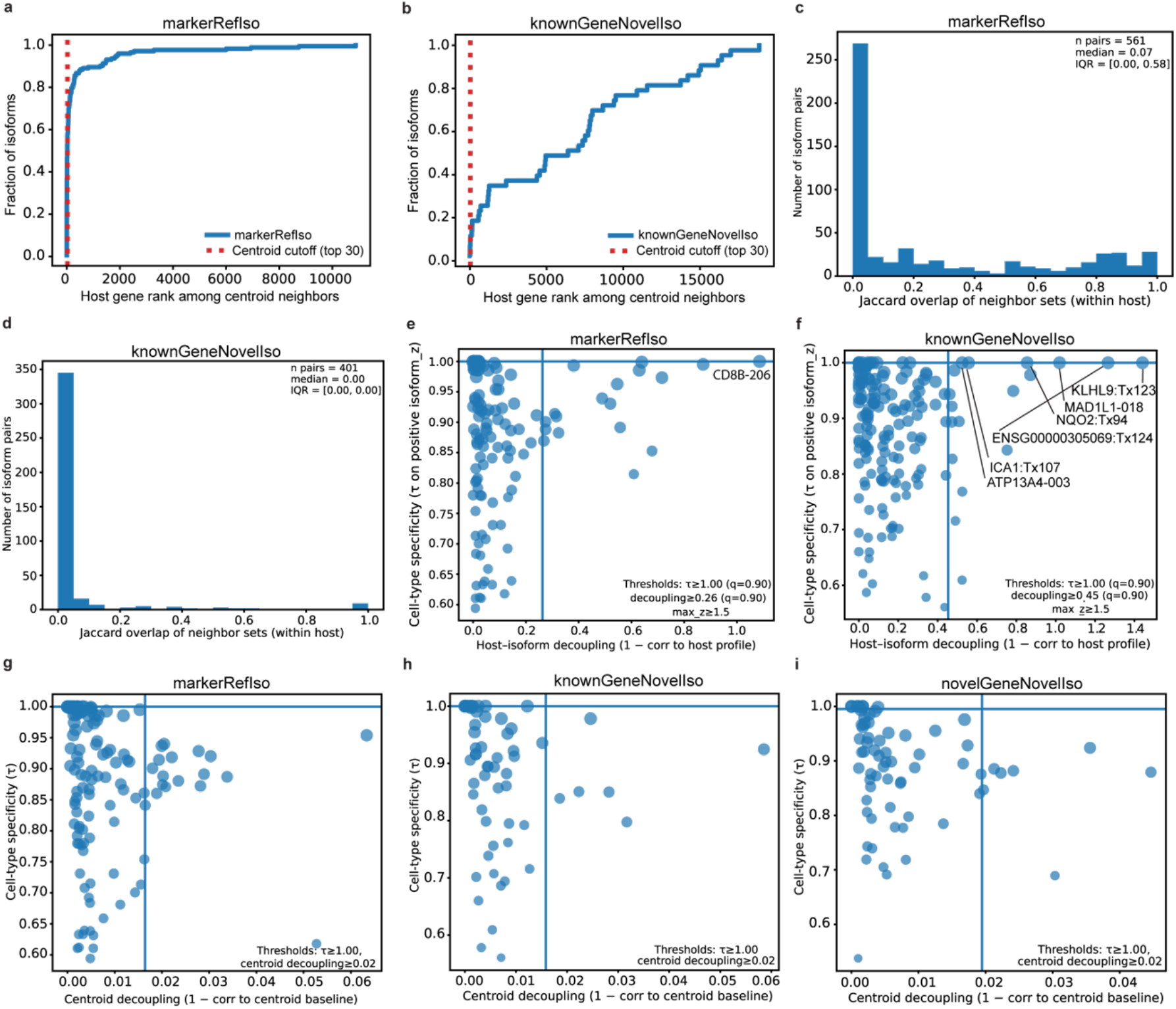
Centroid-based gene reference neighborhoods preserve biologically meaningful expression patterns beyond host gene. **(a-b)** Host gene rank among top most-similar centroid neighbors for each isoform in the a) canonical immune marker reference isoforms and b) novel isoforms from known genes. **(c-d)** Distribution of Jaccard similarity overlap of centroid neighborhoods between isoforms from the same gene for c) *markerRefIso* and d) *knownGeneNovelIso*. Lower Jaccard overlap indicates divergence from isoforms from the same gene. **(e-f)** Isoform decoupling from host gene expression (1-Jaccard overlap) versus cell-type specificity (τ) for (**e**) marker isoforms and (**f**) known genes with novel isoforms. Lines denote 90^th^ percentile (q=0.9), and isoforms above both thresholds were labeled with their gene and transcript name. Novel isoforms are indicated with “Tx” before their isoform number. **(g-i)** Isoform decoupling from centroid neighborhood versus cell-type specificity (τ) for (**g**) *markerRefIso*, (**h**) *knownGeneNovelIso*, and (**i**) *novelGeneNovelIso*. Lines represent the 90^th^ percentile (q=0.9) and isoforms above both thresholds, if any, were labeled with their gene and transcript name.

Among marker isoform neighborhoods, host genes remain enriched within the top 30 most similar centroid neighbors, and are therefore included in the neighborhood, without consistently occupying the first rank, demonstrating that the centroid framework successfully captures biologically relevant neighborhoods while maintaining the unique isoform-level expression diversity. Rather than the isoform centroid neighbors serving as an identity classifier, these groupings reflect functional similarity based on shared cell-type expression patterns, supporting their relevance for downstream analysis of novel isoforms. Interestingly, when this same analysis was performed for neighborhoods belonging to novel isoforms from known genes, not a single isoform had its host gene as the most similar neighbor, and only 1.7% of isoforms had the host gene in the top five centroids (**Figure 2b**). The median rank of host genes among their new isoforms from known genes was 6,392, demonstrating a striking deviation from the host gene cell-type expression profile across the set of reference centroids.

To assess whether isoforms within a known gene (*markerRefIso* and *knownGeneNovelIso)* shared overlap in their centroid neighborhoods, we quantified pairwise neighborhood overlap using Jaccard similarity^21^. The marker reference set showed a fair amount of variance in Jaccard similarity (Interquartile range [IQR] = 0.0-0.58), but the median similarity remained low, at 0.07, indicating very little overlap between neighborhoods (**Figure 2c**). In the dataset of known genes with novel isoforms, we identified very little variance (IQR = 0.0-0.0), with most pairs collecting at 0 and a median of 0.0 (**Figure 2d**). For both marker isoforms and known genes with novel isoforms, an isoform’s neighborhood contained a fairly unique makeup of gene centroids, even among isoforms originating from the same gene.

We sought to determine how isoform expression patterns diverged from their host genes, and whether that difference was cell-type-specific. Among isoforms from cell-type marker genes, a large number of isoforms displayed cell-type specificity, as expected (τ threshold = 1.0), and relatively low decoupling from the host isoform’s expression signature in a cell-type-specific manner (decoupling threshold = 0.26) (**Figure 2e**). Isoforms in this dataset that majorly deviated from their host gene’s expression profile did so in a largely non-cell-type-specific manner, except for CD8B-206 (ENST00000431506), an isoform discussed in this dataset’s original paper due to its unique enrichment in memory T cells compared to CD8*^+^* effector T cells. Among known genes with at least one novel isoform detected, we identified high cell-type specificity and moderate decoupling (τ threshold = 1.0; decoupling threshold = 0.45). Six isoforms surpassed both thresholds with a mixture of both known and novel isoforms deviating in a cell-type-specific manner (**Figure 2f**). By comparison, however, the centroid-based divergence analyses demonstrated high cell-type-specificity (τ threshold = 1.0) with very low centroid decoupling overall (decoupling threshold ≅0.02: *markerRefIso*=0.016, *knownGeneNovelIso*=0.016, *novelGeneNovelIso*=0.019) (**Figure 2g-i**). These analyses reflect higher confidence in the goodness-of-fit of these centroid neighborhoods compared to that of the host gene.

### Reference gene centroids accurately reflect cell-type-specific patterns found in isoforms of interest

In building our centroid neighborhoods, we included the top 30 most similar centroid neighbors with a Pearson similarity r > 0.8. We were interested in how well these neighborhoods built from reference expression profiles fit the assigned isoform. Residuals were calculated to represent how much isoform expression patterns deviated from the baseline expression profiles represented in their centroid neighborhoods. To determine the extent to which our isoforms deviated from their respective neighborhoods, we examined the distribution of the absolute value of maximum residuals for each analysis group. Before these analyses, we set a threshold of |residual| > 1.0 as a conservative cutoff to identify isoforms exhibiting large deviations from centroid neighbor cell-type expression patterns, and to better identify major changes in expression profile.

Among our positive control marker isoforms, the maximum absolute residuals ranged from 0.007 to 0.642, with a median of 0.15 (**Figure 3a**). For novel isoforms from known genes, absolute residuals ranged from 0.009 to 0.678, with a median of 0.25 (**Figure 3b**). Neither of these two groups showed isoforms with |residual| greater than 1.0, indicating these centroid neighbors are closely correlated to their respective isoforms. Among novel isoforms from new genes, maximum |residual| ranged from 0.003 to 1.106 with a median of 0.26. One isoform, *BambuTx111* from *BambuGene58808*, showed a maximum |residual| over 1.0 (**Figure 3c,d**). This deviation, which represents how much the isoform expression pattern deviated from its predicted baseline, was confined to a single cell type, effector-memory transition T cells (|residual|=1.106). This isoform retained overall expression patterns similar to the reference centroid baseline but displayed more cell-type specificity (**Figure 3d**), supporting continued confidence in its centroid neighborhood.

**Figure 3.**
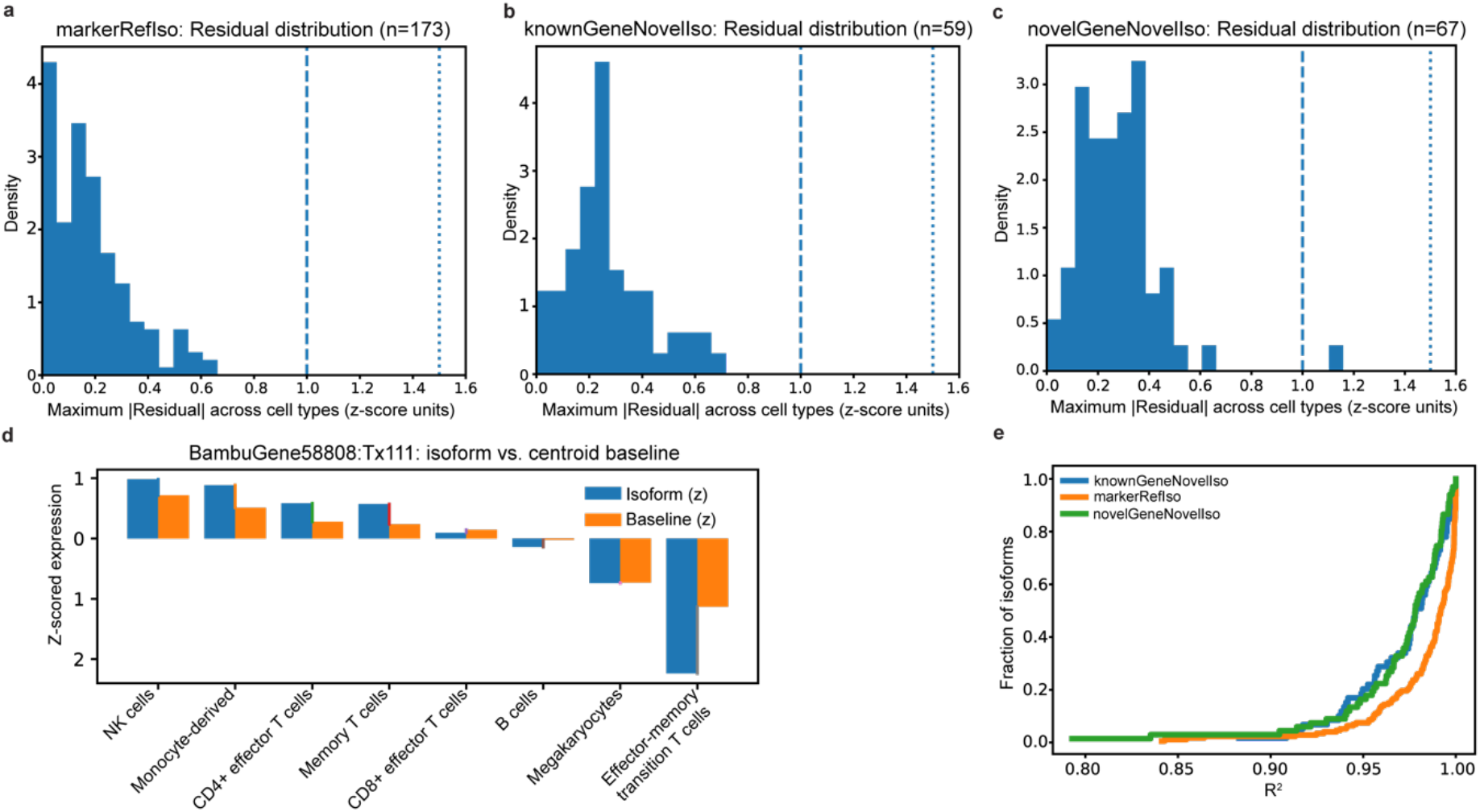
Validation of centroid-based isoform neighborhoods using marker and novel isoforms. **(a-c)** Distributions of maximum absolute residuals across cell types for each isoform, representing deviation between observed isoform expression profile and centroid expression profile, shown for *markerRefIso* (**a**), *knownGeneNovelIso* (**b**), and *novelGeneNovelIso* (**c**). **(d)** Example of isoform-level deviation for *BambuGene58808:BambuTx111* showing observed isoform expression profile (blue) compared to baseline predicted expression profile according to centroid baseline (orange) across immune cell types. **(e)** Goodness-of-fit analysis between isoforms and their corresponding centroid neighborhoods, shown as distributions of the coefficient of determination (R^2^) across isoforms in each analysis set.

Each centroid neighborhood was highly similar to its isoform of interest, with minimum Pearson correlation coefficients of 0.937 for *markerRefIso*, 0.941 for *knownGeneNovelIso*, and 0.955 for *novelGeneNovelIso*. Consistent with the high Jaccard similarity shown in these neighborhoods, the reference centroid baselines represented the majority of the isoform-level expression variance across cell types (**Figure 3e**). Together, these results indicate that these centroid neighborhoods robustly capture a structure that faithfully represents each isoform’s expression pattern profile, supporting their use as a reliable reference for downstream analyses.

### Evaluation of centroid neighborhoods by pathway enrichment, using cell-type marker isoforms as a positive control

To evaluate whether the centroid neighborhood framework could identify biologically meaningful pathway associations for isoforms of interest, we performed pathway enrichment for our marker isoforms using KEGG^36^, GO^28,29^, and Reactome^37,40^. We used these three annotation sources to identify consistent biological themes while minimizing bias from using just one. The top 12 significant pathways per source (by number of isoform centroid neighborhoods enriched) were plotted in **Figure 4**. As expected, centroid reference genes derived from immune cell marker isoforms largely reflected immune-related biological processes across the three databases (**Figure 4**).

**Figure 4.**
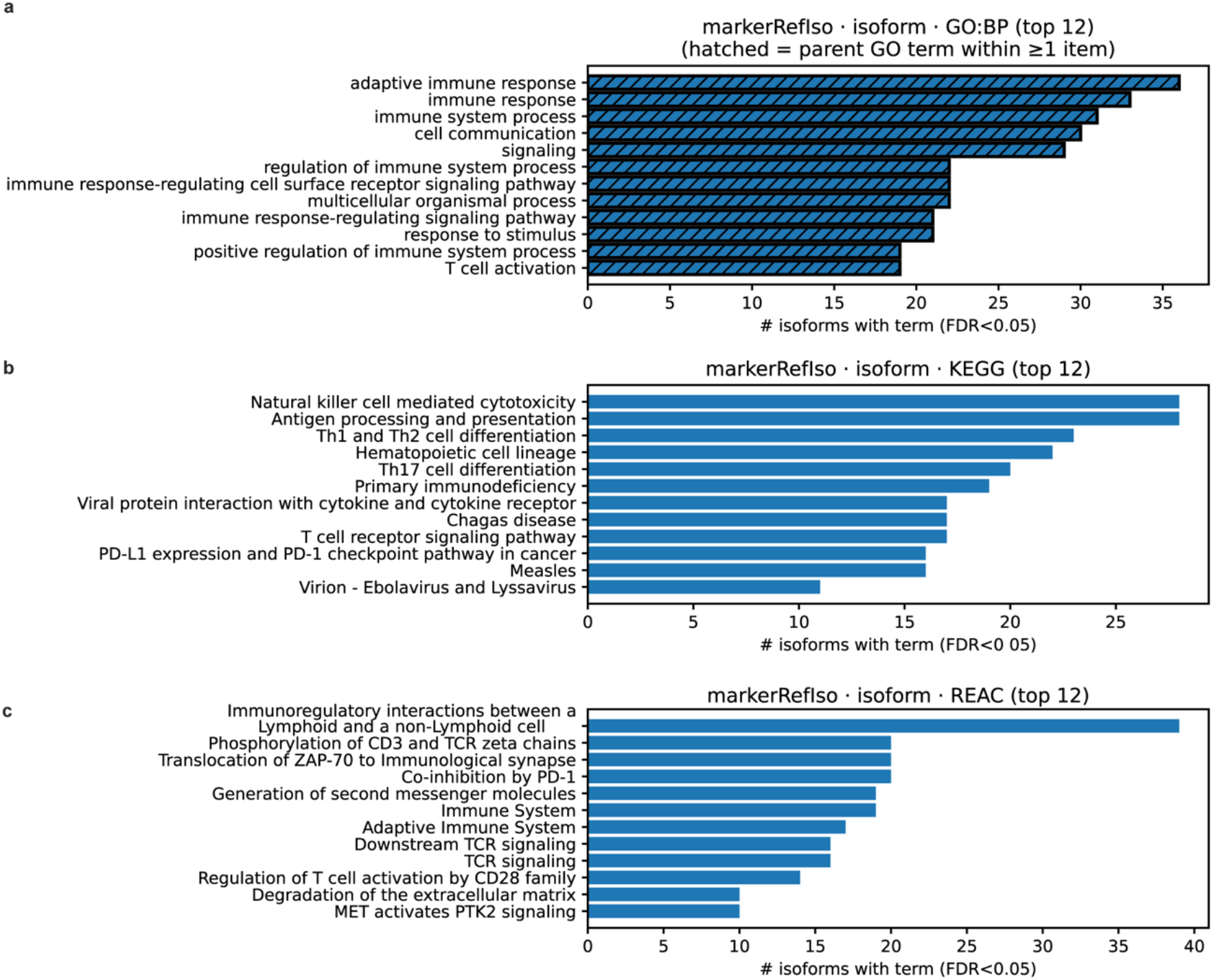
Validation of centroid-based isoform neighborhoods using pathway analysis strategies on marker isoforms. **(a-c)** Overall pathway enrichment of centroid neighborhood genes associated with reference marker isoforms. Annotation sets used for this figure include Gene Ontology Biological Processes (GO:BP, **a**), Kyoto Encyclopedia of Genes and Genomes (KEGG, **b**), and Reactome pathway database (REAC, **c**). For GO-based analyses, “parent” biological process terms that encompass at least one other specific “child” term were indicated with a hatched bar to indicate their status as such. Bars represent the top 12 pathways by number of isoforms associated with each significantly enriched item (FDR < 0.05), then FDR from lowest to highest.

For the GO-based figures, we represented the hierarchical structure of the ontology by reporting the parent pathways that encompassed at least 1 more specific “child” pathway underneath. GO enrichment included pathways composed solely of parent ones related to broad immune response pathways (**Figure 4a**). The top GO pathway was “adaptive immune response” (GO:0002250), which was not unexpected given that T cell subtypes and B cells were highly prominent in our dataset. By comparison, KEGG pathways showed more specificity, exhibiting a mixture of cell-type-related pathways (e.g., Natural killer cell mediated cytotoxicity (hsa04650), Th1 and Th2 cell differentiation (hsa04658), Hematopoietic cell lineage (hsa04640), and viral or parasitic response pathways (e.g., Measles (hsa05162), Chagas disease (hsa051142) (**Figure 4b**). Reactome pathways represented a mixture of broad and specific terms, with the most centroid neighborhoods enriched for “Immunoregulatory interactions between a Lymphoid and a non-Lymphoid cell” (R-HSA-198933)^42^ (**Figure 4c**). The biological themes shared between GO and Reactome related to cellular communication between cell types, immune response (specifically highlighting adaptive), and enrichment of T cell-specific pathways, whereas a common theme between the Reactome and KEGG pathways included T cell receptor signaling.

To validate that our cell-type labels were supported by expected pathway-level biological signatures, we split our centroid-derived gene sets by cell-type marker and performed pathway enrichment analysis using GO, KEGG, and Reactome annotations (**Figure 5**). Across the top 12 pathways from all three annotation sets, T cells and natural killer (NK) cells both showed enrichment for multiple pathways from the GO, Reactome, and KEGG annotation sets, whereas B cells were more prominently represented by the Reactome annotations, and monocyte-derived cells and megakaryocytes were more enriched for GO Biological Process (GO:BP) terms. T cells, NK cells, and monocyte-derived cells all demonstrated enrichment of broad terms related to immune function (e.g., adaptive immune response, GO:0002250) within the top twelve terms, while B cells and megakaryocytes did not. NK cells and monocyte-derived cells also exhibited enrichment for other broad terms, including cell communication, signaling, or regulation of biological processes.

**Figure 5.**
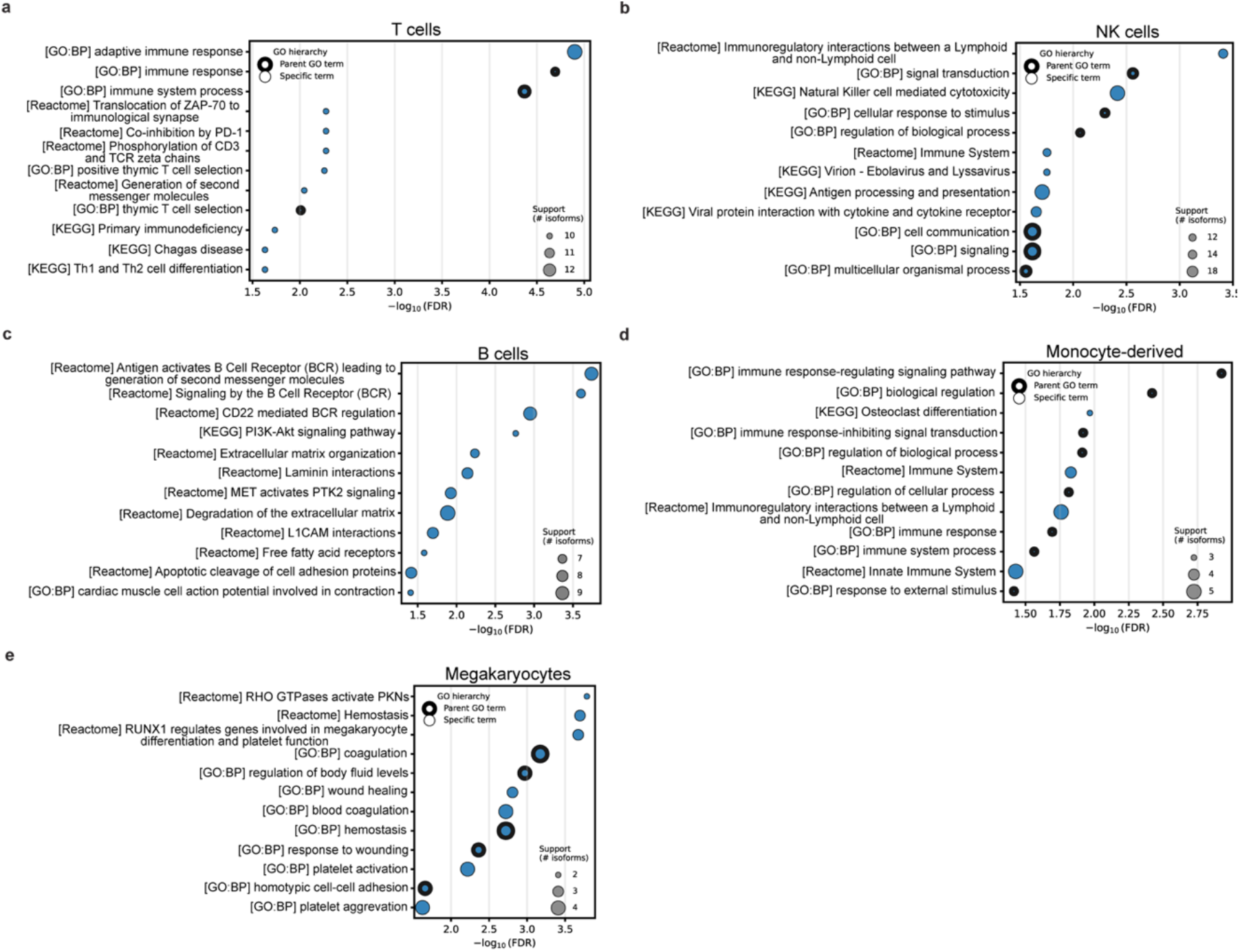
Pathway analyses of genes from centroid neighborhoods belonging to cell-type marker gene isoforms reflect biological functions consistent with the cell-type of interest. **(a-e)** Pathway enrichment of genes from centroid neighborhoods associated with reference marker isoforms. Enrichment was performed using Gene Ontology Biological Process (GO:BP), Kyoto Encyclopedia of Genes and Genomes (KEGG), and Reactome databases. The top 12 pathways across databases (as determined by lowest FDR) are split by cell-type grouping for markers including **(a)** T cells—*CD3D*, *CD3E*, *CD3G*, **(b)** NK cells—*GZMB*, *KLRF1*, *NCAM1*, *ITGAM*, *IL2RB*, **(c)** B cells—*CD22*, *CD79A*, *MS4A1*, *CD19*, **(d)** Monocyte-derived cells—*FCGR2A*, *CLEC7A*, *CD33*, *LILRB4*, **(e)** Megakaryocytes—*GP1BA*, *MPL*, *ITGA2B*. For GO-based analyses, any biological process term that encompassed at least one or more specific child terms was outlined in black. The bubble size indicates the number of isoform centroid neighborhoods supporting each enriched pathway.

T cells are major mediators of adaptive immunity and are typically split into CD4^+^ (helper) or CD8^+^ (cytotoxic) subpopulations. T cells show tremendous functional diversity, demonstrating phenotypes with roles in immune system regulation, cytotoxicity, and cytokine production and secretion^43^, and differentiate into specialized subtypes upon activation according to the cytokine responsible for their activation^44^. T cell pathway enrichments across all three annotation sets (**Figure 5a**) were consistent with processes specific to these immune cells including thymic T cell selection (GO:0045061), Th1 and Th2 cell differentiation (hsa04658), and phosphorylation of CD3 and TCR zeta chains (R-HSA-202427) Several disease-associated pathways appearing in the top enriched terms reflect immune processes T cells play a major role in (e.g., Chagas disease (hsa05142)^45^, Translocation of ZAP-70 to immunological synapse (R-HSA-202430)^46^, Co-inhibition by PD-1 (R-HAS-389948)^47^), making their enrichment expected.

NK cells are innate lymphoid cells that participate in the rapid, targeted apoptosis of cells and immune surveillance, important for removing infected or malignant cells from the body^48,49^. They can also assist in cytokine production that can shape broader immune responses^48,49^. Neighbors for NK cell-associated marker isoforms (**Figure 5b**) were enriched for pathways consistent with their known functionality, including antigen processing and presentation (hsa04612), and natural killer cell mediated cytotoxicity (hsa04650). Multiple pathways associated with immune responses to viral infection were also among the top 12 pathways, consistent with the role of these cells in innate antiviral response^50^.

B cells are also key players of adaptive immunity and are responsible for antibody and cytokine production as well as antigen presentation^51^. Pathways tied to B cell marker isoform neighbors (**Figure 5c**) were most enriched for terms associated with B cell receptor signaling or B cell-specific functions (e.g., CD22 mediated BCR regulation, R-HSA-5690714^52^) Several pathways related to extracellular matrix and adhesion signaling also appear, including extracellular matrix organization (R-HSA-1474244), laminin interactions (R-HSA-30000157)^53^, and MET activated PTK2 signaling (R-HSA-8874081). These pathways may reflect B-cell migration and adhesion programs that interact with B cell receptor signaling pathways, including inhibitory regulation via CD22 during B cell migration throughout the body^427–429^. Enrichment of free fatty acid receptors (R-HSA-444209) may reflect metabolic changes that occur in B cell activation, which have shown direct relevance to autoimmune disorders, including Multiple Sclerosis (MS)^54,55^. The most surprising pathway tied to B cell markers is the “cardiac muscle cell action potential involved in contraction” Reactome pathway (R-HSA-5576891), since its title indicates a highly cardiac-specific function. However, the similarities in calcium influx during an action potential and B cell receptor signaling^56^ likely explain this connection, rather than reflecting actual cardiac-specific biology.

Monocytes display functions associated with patrolling for pathogenic behavior, phagocytosis, antigen presentation, and cytokine production^57^. Monocyte-derived cells in our dataset likely encompass a heterogeneous range of myeloid lineage cells, including classical, non-classical, and plasmacytoid dendritic cells^58^, as well as monocytes themselves, based on our markers. These cells act as circulating innate immune sensors and regulators, responding to stimuli and differentiating into different cell types as necessary^58,59^. This may explain the broad representation of terms in the top 12 pathways enriched for gene neighbors associated with isoform markers of this cell type (**Figure 5d**). One exception is the “Osteoclast differentiation” KEGG pathway (hsa04380), in which monocyte/macrophage precursor cells develop into specialized bone reabsorption cells, osteoclasts^60^.

Lastly, megakaryocytes are known for their role in platelet production, but have also been more recently recognized as being highly diverse functionally^61–68^, appearing to play a role in platelet generation, immune response, hematopoiesis, and immune surveillance. Neighbors for megakaryocyte-associated markers were strongly enriched for pathways related to platelet production and function (**Figure 5e**), including coagulation (GO:0050817, GO:007596), hemostasis (GO:0007599), and cell aggregation (GO:0098743). The top pathway is the Reactome-based “RHO GTPases activate PKNs” (R-HSA-5625740), which supports cytoskeletal remodeling programs associated with megakaryocyte maturation, since RHO GTPases are major regulators of megakaryocyte development, differentiation, and platelet production^69^. Multiple pathways enriched in megakaryocytes support active megakaryocyte differentiation and platelet production.

Pathway enrichment results were consistent with each cell-type’s established roles in the immune system, providing support for the assigned cell-type labels as well as the biological relevance of the centroid neighborhoods. These results indicate that centroid neighborhoods capture biologically meaningful expression patterns, and can provide useful pathway-level context for isoforms, as shown with known isoforms from established cellular marker genes. Notably, centroid-based references retained biologically relevant signals of isoforms not found on the gene level alone.

Our goal was to use cell-type expression patterns to infer pathway context for new isoforms, so we next plotted the top 30 novel isoforms by cell-type specificity from known genes and from previously unannotated genes. The cell types with the greatest number of new isoforms from known genes that were cell-type-specific (**Figure 6a**) were B cells, T cells, and megakaryocytes, each exhibiting enrichment of seven new isoforms. Among isoforms from previously unannotated genes (**Figure 6b**), the greatest number of isoforms were identified in monocyte-derived cells, followed closely by NK cells, with nine and eight isoforms enriched, respectively.

**Figure 6.**
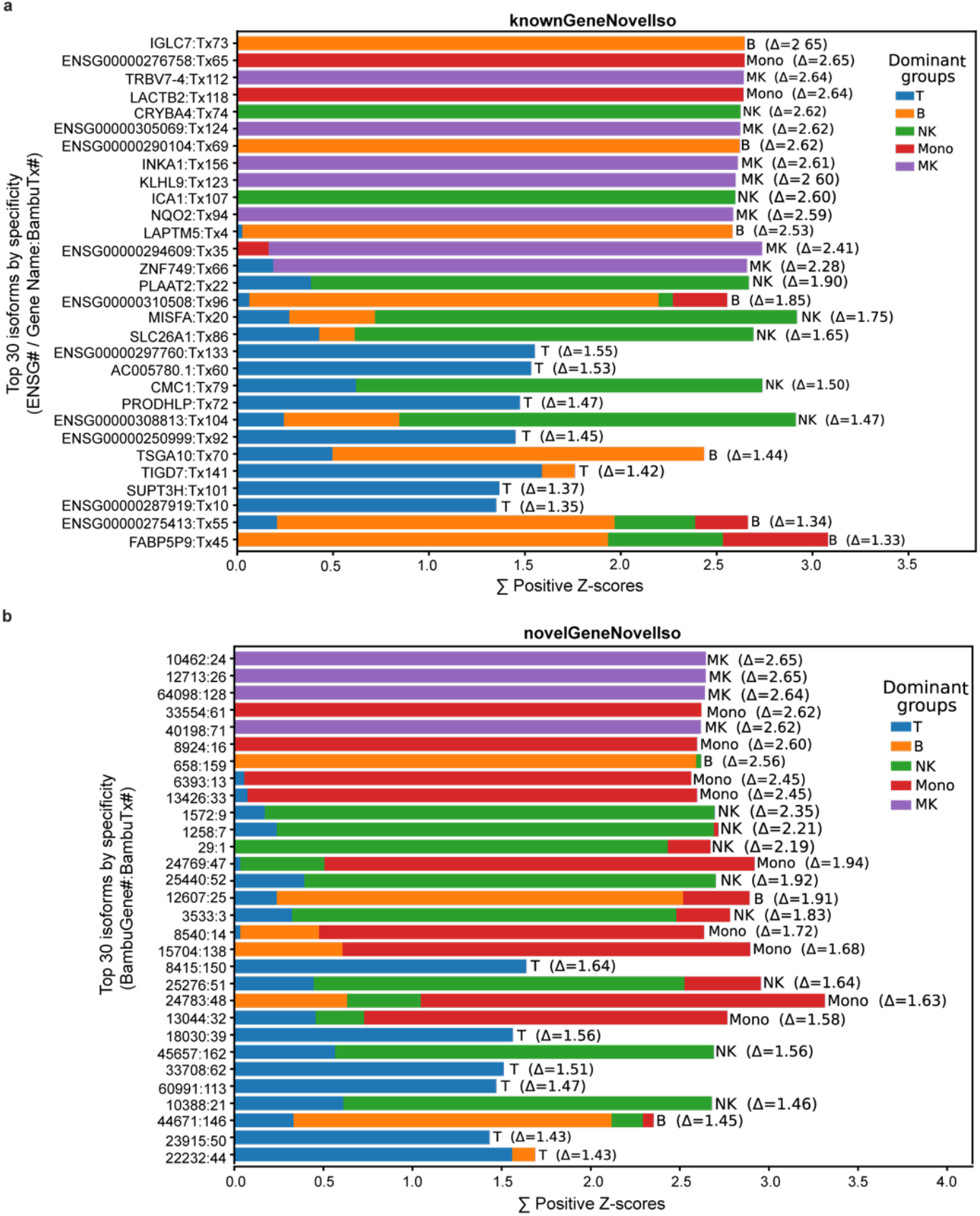
Cell-type enrichment of novel isoforms from previously annotated and unannotated genes. **(a-b)** Cell-type enrichment for the top 30 most cell-type-specific novel isoforms from a) known and b) new genes, ranked by cell-type specificity score (Δ). Bars represent major immune cell group enrichment, summarizing positive isoform z-scores across all major immune cell groups (T cells, NK cells, B cells, monocyte-derived cells, megakaryocytes), with the height of the bar representing the sum (Σ) of positive isoform z-scores within an individual isoform. Colored segments within the stacked bar indicate individual cell-type z-scores contributing to positive enrichment within the group. A dominant immune cell was assigned for each isoform based on the highest cell-type specificity score (Δ), where group scores were evaluated as the mean of the two highest positive isoform z-scores within a single major cell type.

For new isoforms from known genes, we also compared the reported cell-type-specific patterns from the EMBL-EBI Expression Atlas^70^ and Human Protein Atlas^71–75^ to identify a consensus in the specificity between these two sources. Five of our known genes with a novel isoform labeled as long non-coding RNA (lncRNA) genes according to GeneCards (V5.26)^76^ did not have reported cell-type expression data in both datasets, and could not be included in our comparison analysis. Specifically, these genes were: *ENSG00000287919*, *ENSG00000308813*, *ENSG00000297760*, *ENSG00000305069*, and *ENSG00000310508*. *ENSG00000287919* is the only one of these five genes found in either database, and was reported as being detected in the cerebellar hemisphere of the GTEx dataset^77^, a dataset produced via bulk long-read sequencing. All these genes are listed as novel genes or transcripts in the GeneCards database (therefore, newly discovered by another study)^76^. *ENSG00000305069* is the only one of these known genes with a relationship to an existing gene and is listed as being sense intronic to *TLN1*^76^. In our dataset, we identified two of these genes as demonstrating T cell enrichment (*ENSG00000287919* and *ENSG00000297760*) and the other three are enriched for other cell types (**Figure 6a**). The rarity of these genes in published datasets may relate to the difficulty in detecting them with short-read or bulk methodologies.

11 of our 25 new isoforms from known genes included in this analysis exhibited high expression patterns in the same cell type or in its precursor, reported at the gene level in the EMBL-EBI Expression Atlas and/or Human Protein Atlas (*IGLC7*, *TRBV7-4*, *INKA1*, *ICA1*, *NQO2*, *LAPTM5*, *ZNF749*, *CMC1*, *TIGD7*, *SUPT3H, ENSG00000276758*, *ENSG00000290104*). An additional four genes reported relatively high expression in the cell type enriched by our new isoform but were not the most highly enriched cell type according to previously reported gene-level data (*CRYBA4*, *ICA1*, *SLC26A1*, *AC005780*). Eight of our known genes with new isoforms did not show cell-type enrichment in either of our reference gene datasets, instead exhibiting broad expression across multiple cell types without a standout immune cell-type of interest (*KLHL9*, *LACTB2*, *TSGA10*, *PRODHLP*, *SLC26A1*, *MISFA*, *ENSG00000250999*, *ENSG00000275413*). Some of these, instead, were more enriched in other sources rather than being most highly expressed in blood-based immune cells. For example, *SLC26A1* and *KLHL9* were both more highly expressed in mesenchymal cells, whereas *MISFA*, *TSGA10*, *ICA1*, and *LACTB2* were most common in spermatogenic cell types. Importantly, however, expression is not exclusive to these areas, and their apparent specificity may reflect limitations of capturing low-abundance or alternative transcripts using short-read or bulk methodologies, and a need to better understand the role of these genes in peripheral immune cells.

### Pathway analyses of centroid neighbors for novel isoforms from known genes suggest associations with transcriptional and translational regulation

We next performed pathway enrichment for the genes associated with new isoforms from previously annotated genes via their centroid neighborhoods. As in our positive control analyses using our known marker gene isoforms, we used GO:BP, KEGG, and Reactome for our pathway annotation sets in order to minimize bias from using only one annotation source. Our objective with this analysis was to identify common signatures tying together these unknown isoforms that can be further dissected.

Among the GO:BP terms (**Figure 7a**), the most heavily enriched terms were dominated by a direct connection to RNA production. The top two GO:BP terms (GO:0006228, GO:0046051) centered on UTP, uridine triphosphate, a crucial ribonucleoside triphosphate that is a major building block of RNA as well as a source of energy in processes such as glycogen synthesis^78^. The other eight terms describe the synthesis and metabolism of specific components of RNA building blocks, such as pyrimidine nucleoside triphosphate. Within the context of PBMCs, the inclusion of the “anatomical structure morphogenesis” pathway (GO:0009653) may reflect immune cell migration and participation in tissue repair and remodeling after injury^79,80^, rather than directly related to developmental processes. Similarly, the “sensory organ development” pathway (GO:0007423) could be related to cell signaling and calcium signaling required for cellular activation in response to immune threats, as shown above and in **Figure 5**.

**Figure 7.**
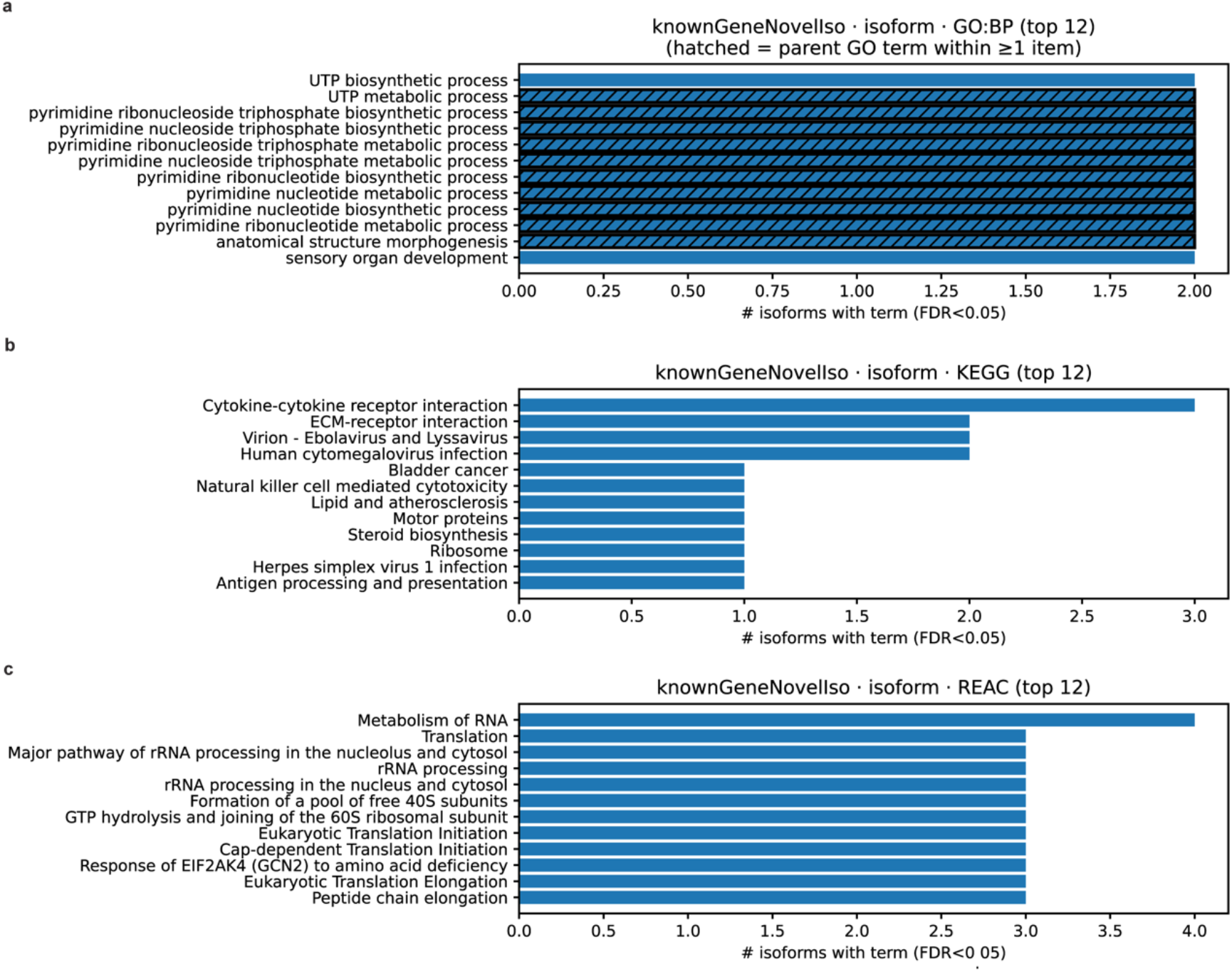
Overview of pathways enriched in new isoforms from known genes. **(a-c)** Overall enrichment of the top 12 pathways called by centroid neighbor genes tied to novel isoforms from known genes. Annotation sources used for this figure include Gene Ontology: Biological Processes (GO:BP, **a**), Kyoto Encyclopedia of Genes and Genomes (KEGG, **b**), and Reactome pathway database (REAC, **c**). GO-based analysis included highlighting “parent” biological process terms encompassing at least 1 other “child” term using a hatched bar to indicate their status. Pathway ranking was determined according to the number of isoforms associated with each significantly enriched item (FDR < 0.05), then sorted from lowest FDR to highest if multiple terms for the same isoform number were detected.

KEGG pathway enrichment (**Figure 7b**) showed a greater representation of immune-related pathways than in the GO:BP analysis. The top pathway, “cytokine-cytokine interactions” (hsa04060), as well as other top pathways related to viral infection (hsadd03265, hsa05163, hsa05168), suggest that some of these novel isoforms from known genes may be associated with expression patterns linked to immune activation in infection or disease. This interpretation is also supported by pathways tied to antigen presentation (hsa04612) and cytotoxicity response from natural killer cells (hsa04650). The “bladder cancer” pathway (hsa05219) includes a large number of immune signaling molecules^81^, which may explain its enrichment here independent of cancer-specific biology. Lastly, KEGG also represented housekeeping pathways, including “Ribosome” (hsa03010) and “ECM-receptor interaction” (hsa04512).

The top Reactome pathways (**Figure 7c**) were consistent with enrichment for RNA processing and protein synthesis pathways. The “Metabolism of RNA” superpathway (R-HSA-8953854) was highly enriched, composed of multiple participating events, including three highlighted in the top 12 terms (rRNA processing, R-HSA-72312; rRNA processing in the nucleus and cytosol, R-HSA-8868773; Major pathway of rRNA processing in the nucleolus and cytosol, R-HSA-6791226). The “Metabolism of RNA” superpathway is associated with the “RNA metabolic process” (GO:0006364) pathway, but in our dataset, represents more isoform centroid neighborhoods than the GO:BP event. The Reactome “Translation” pathway (R-HSA-72766), the second most highly supported Reactome term (**Figure 7c**), also contains more specific events; among them are six of our top Reactome pathways (Peptide chain elongation, R-HSA-156902; Eukaryotic translation elongation, R-HSA-156842; Response of EIF2AK4 (GCN2) to amino acid deficiency, R-HSA-9633012; Cap-dependent translation initiation, R-HSA-72737; GTP hydrolysis and joining of the 60S ribosomal subunit, R-HSA-72706; Formation of a pool of free 40S subunits, R-HSA-72689).

Together, these pathway analyses indicate a subset of novel isoforms from known genes is associated with expression patterns tied to transcriptional and translational regulatory processes, particularly RNA production and metabolism and translation-related programs. These findings support the notion that our centroid neighborhoods can capture biologically meaningful pathway-level signals for novel isoforms across multiple cell types, rather than only for cell-type-enriched marker isoforms alone.

### Pathway analyses of centroid-neighbor gene sets suggest diverse pathway associations for novel isoforms from new genes

To further test the capabilities of combining centroid neighborhoods with pathway enrichment to provide pathway-level context for novel isoforms, we used our set of novel isoforms from previously unannotated genes (**Figure 8**). In the absence of a known host gene, these isoforms are truly untethered from established functional annotations and subsequent assumed functionality, so these analyses represent a stringent test for our computational framework. As observed with novel isoforms from known genes, fewer isoform neighborhoods were enriched for any single term compared to marker isoforms. The number of new isoforms from previously unannotated genes per enriched term was similar to those from known genes, with no single pathway supported by more than five isoform neighborhoods.

**Figure 8.**
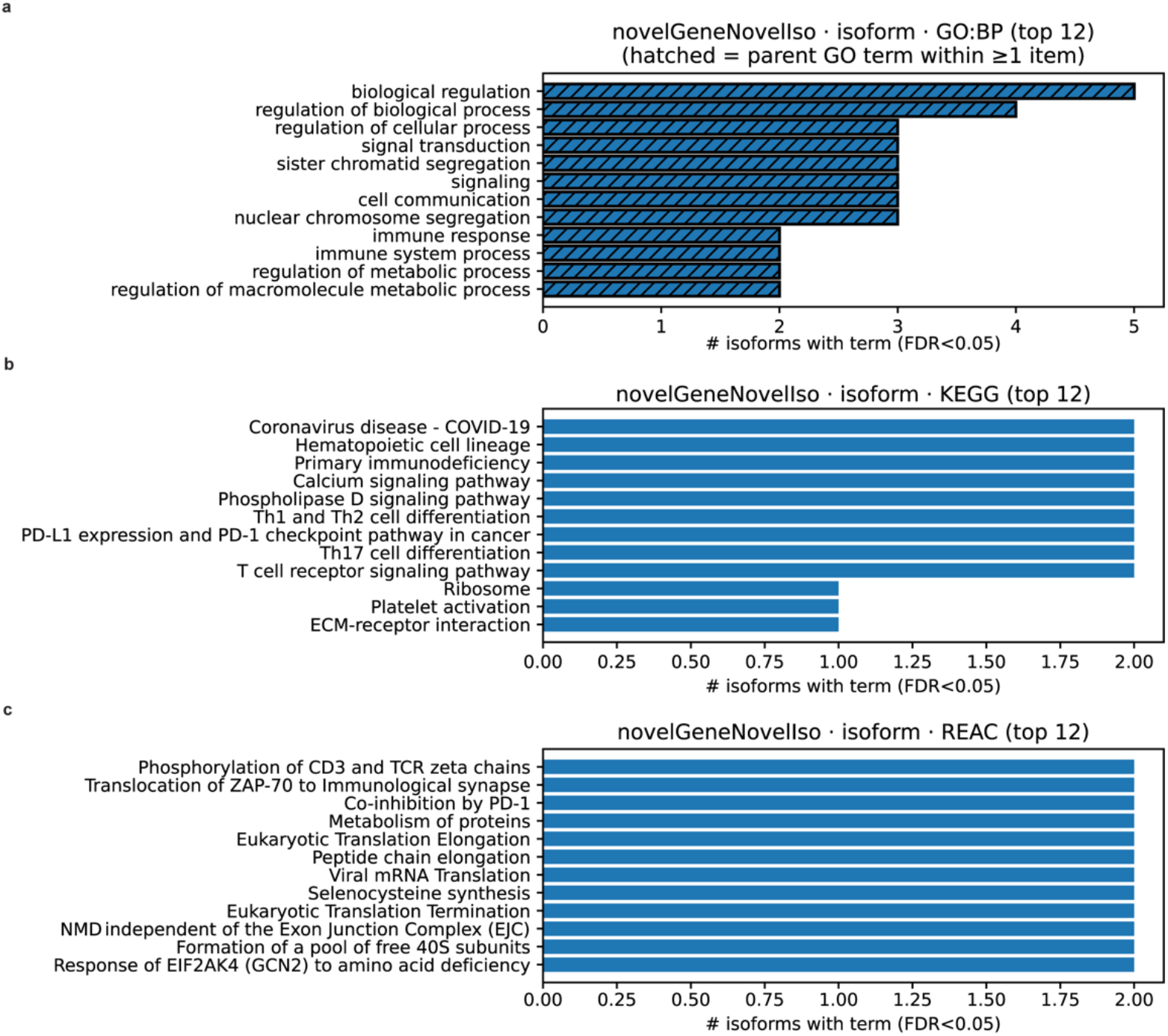
Overview of pathways enriched in new isoforms from new genes. **(a-b)** Top 12 biological pathways enriched by centroid neighbor genes associated with new isoforms from new genes. Annotation sources for pathway enrichment included **a)** Gene Ontology: Biological Processes (GO:BP), **b)** Kyoto Encyclopedia of Genes and Genomes (KEGG), **c)** Reactome pathway (REAC). GO:BP analyses highlight broader parent terms that include at least one other significant “child” term using a hatched bar. Pathway rankings are assessed according to the number of isoforms associated with terms with an FDR <0.05.

The GO:BP terms represented by the novel isoform neighborhoods (**Figure 8a**) were largely broad biological processes. The top four terms represent a cascade from broad parent terms to more specific child ones, likely reflecting enrichment driven by the same isoforms. Specifically, “biological regulation” (GO:0065007) is a parent term of “regulation of biological process” (GO:0050789), which is a parent term of “cellular process” (GO:0009987), a parent term of “signal transduction” (GO:007165). Unlike our new isoforms from known genes, GO enrichment in this isoform set included immune-specific pathways, indicating that a portion of our new isoforms from previously unannotated genes may be associated with immune regulatory pathway signatures. The enrichment of metabolic pathways (GO:001922, GO:0060255) may also relate to immune regulation, since metabolic programming is known to shift under immune activation^82,83^. The pathways associated with the segregation of nuclear chromosomes (GO:0098813) and sister chromatids (GO:000819) may correspond to proliferation-associated programs accompanying immune activation.

The enriched KEGG terms (**Figure 8b**) reflected a more specific aspect of immune response than our GO:BP analyses. The five most highly enriched terms include core immune pathways shared across multiple subtypes of PBMCs, encompassing cellular activation, replication, and immune signaling. Four enriched pathways describe more specific immune processes, including T cell subtype-specific differentiation (hsa04658, hsa04659), signaling in response to cancer (hsa05235) and other immune challenges. “Platelet activation” (hsa04611) was also enriched in one of the isoform neighborhoods, which may be compatible with megakaryocyte-associated immune responses to tissue damage^84^. Lastly, the two remaining pathways reflect housekeeping processing, including ribosomal maintenance (hsa03010) and extracellular matrix-receptor interactions (hsa04512).

Reactome pathways enriched in our novel-isoform-novel-gene neighborhoods (**Figure 8c**) were consistent with the immune signaling themes observed in GO and KEGG analyses. The top three pathways overlapped with terms enriched in T cell marker isoform neighborhoods, including signaling through the T cell receptor complex (R-HSA-202427, R-HSA-202430, R-HSA-389948). The enrichment of selenocysteine synthesis (R-HSA-2408557) may reflect oxidative stress handling in immune cells during activation, or other adaptive immune responses^329–331^. The remaining enriched Reactome terms included multiple translation-related pathways of both viral and eukaryotic origin, suggesting association of a portion of the new isoforms with pathway signatures related to protein synthesis and translational control.

Overall, the newly identified transcripts from previously unannotated genes appear to predominantly exhibit signatures associated with immune-related pathways. However, we see less convergence on a single central function in the new genes than in the known ones, demonstrating greater diversity in our inferred pathway contexts. The presence of pathways tied to immune signaling, metabolic and translational regulation, and housekeeping functionality supports the ability of our centroid framework to infer relevant biologically meaningful pathways, even among diverse novel transcripts.

### Cellular component GO analyses support pathway inference of novel isoforms from biological process enrichment analyses

Given the enriched pathways in our novel isoform neighborhoods related to transcriptional and translational regulation and immune signaling, we sought to evaluate whether inferred functional themes were consistent with expectations for cellular localization. For example, genes involved with translational regulation are typically localized to ribosomes and cytoplasm, while genes associated with T cell receptor signaling are expected to localize to receptor complexes in the plasma membrane. In the context of our framework, an isoform’s centroid neighbors can be used to identify whether inferred functions align with known cellular component annotations. We therefore performed pathway enrichment on the gene-level centroid neighbors of our novel isoform sets using Gene Ontology: Cellular Compartment (GO:CC), displaying up to the top 20 significant terms (FDR<0.05) identified in each isoform group (**Figure 9**).

**Figure 9.**
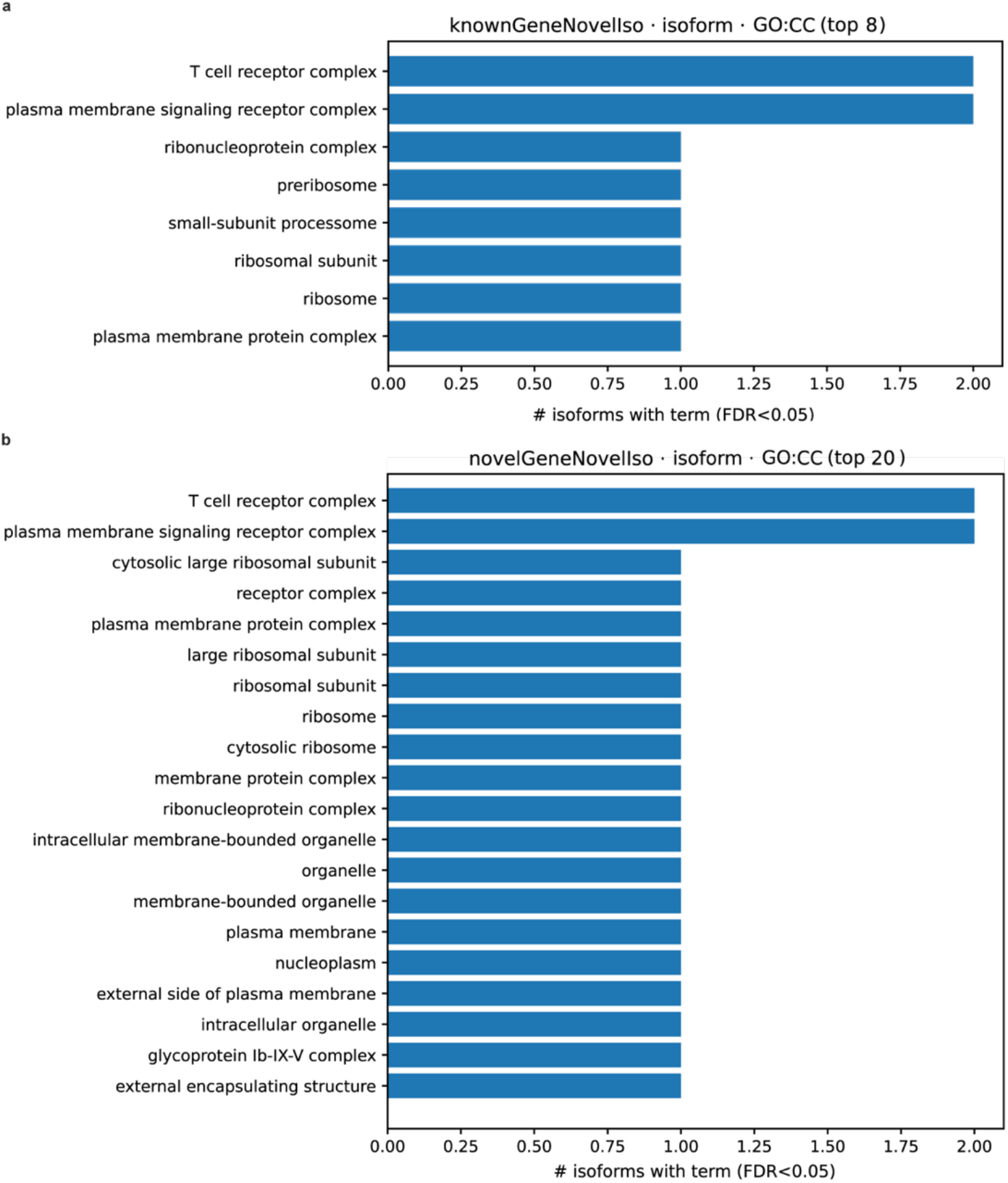
Gene Ontology Cellular Component (GO:CC) pathway annotations. **(a-b)** Up to the top 20 cellular components enriched in centroid gene neighborhoods associated with new isoforms from a) known and b) previously unannotated genes. Annotation source for these terms was from Gene Ontology Cellular Component (GO:CC). Terms with an FDR <0.05 are ranked according to the number of isoform neighborhoods associated with a specific cellular component.

Among terms enriched in centroid neighbors associated with novel isoforms from known genes, only eight terms were deemed to be significant (**Figure 9a**). For this subset of isoforms, we expected enrichment in the cytosol, nucleus, and ribosomes, given the top enrichment of transcriptional and translational processes, as shown in Figure 7. Consistent with this expectation, ribosomes as well as ribosome-adjacent components were enriched, which supports our assertion that a subset of our novel isoforms from annotated genes are involved in translational processes. Notably, the top two compartments reflected cell surface receptor signaling involvement, specifically in the T cell receptor complex, indicating that immune signaling was also highly associated with the inferred functional neighborhoods.

As observed in our GO:BP, KEGG, and Reactome pathway analyses, cellular component terms enriched among centroid neighbors of new isoforms from previously unannotated genes demonstrate a wider array of term diversity than our other novel isoform set, displaying all twenty significant terms (**Figure 9b**). The functional themes shown in our pathway analyses from this isoform set included translational regulation, metabolic processes, and immune signaling, which we would expect to be localized to the nucleus, mitochondria, cytosol, and plasma membrane. However, localization of metabolic processes and immune signaling within the cell can vary depending on specific pathway components involved^85^. As in our other novel isoform dataset, we identified the top cellular compartments to be the T cell receptor complex (GO:0042101) and plasma membrane signaling receptor complex (GO:0098802), suggesting that localization of a portion of immune signaling may be confined to these compartments. We noticed several terms related to ribosomal localization and sublocalization, indicating some amount of translational intervention. Our hypothesis that we would identify mitochondrial and nuclear localization is also supported, due to the enrichment of membrane-bounded organelle and nucleoplasm terms. We also identified enrichment of the glycoprotein Ib-IX-V complex term (GO:1990779), a membrane receptor complex that appears in megakaryocytes and platelets^86^, which is likely tied to the single novel isoform we identified enriched for the Platelet activation KEGG pathway (hsa04611).

Overall, our cellular compartment analyses support the conclusions formed from our pathway enrichment analyses. We expect that based on the similarity of expression profiles and the resultant centroid neighborhoods, that we can infer our novel isoforms to be involved in immune signaling, metabolic processes in the mitochondria, transcription, and translation. From these results, we can begin to narrow down and dissect the function of individual novel isoforms.

### T cell receptor signaling complex localization as a case study for pathway inference of novel isoforms from centroid-enabled pathway analyses

While these pathway enrichment analyses allow inference of trends among isoform sets, they do not illustrate the individual pathway context of our novel isoforms. The top component term being enriched for the T cell receptor (TCR) complex in both of our novel isoform datasets led us to investigate the roles of our novel isoforms whose centroid neighborhoods were associated with this cellular component. In both datasets, we isolated novel isoforms and their centroid neighbor genes associated with the TCR complex and identified two novel genes and four known genes (*PRODHLP*, *SUPT3H*, *ENSG00000267320*, and *ENSG00000297760*) tied to this cellular component term. The relationship between these novel isoforms and their centroid neighbors (**Figure 10**), included multiple TRAV and TRBV protein family members, which are associated with the adaptive aspect of TCR signaling essential for antigen recognition^87–89^.

**Figure 10.**
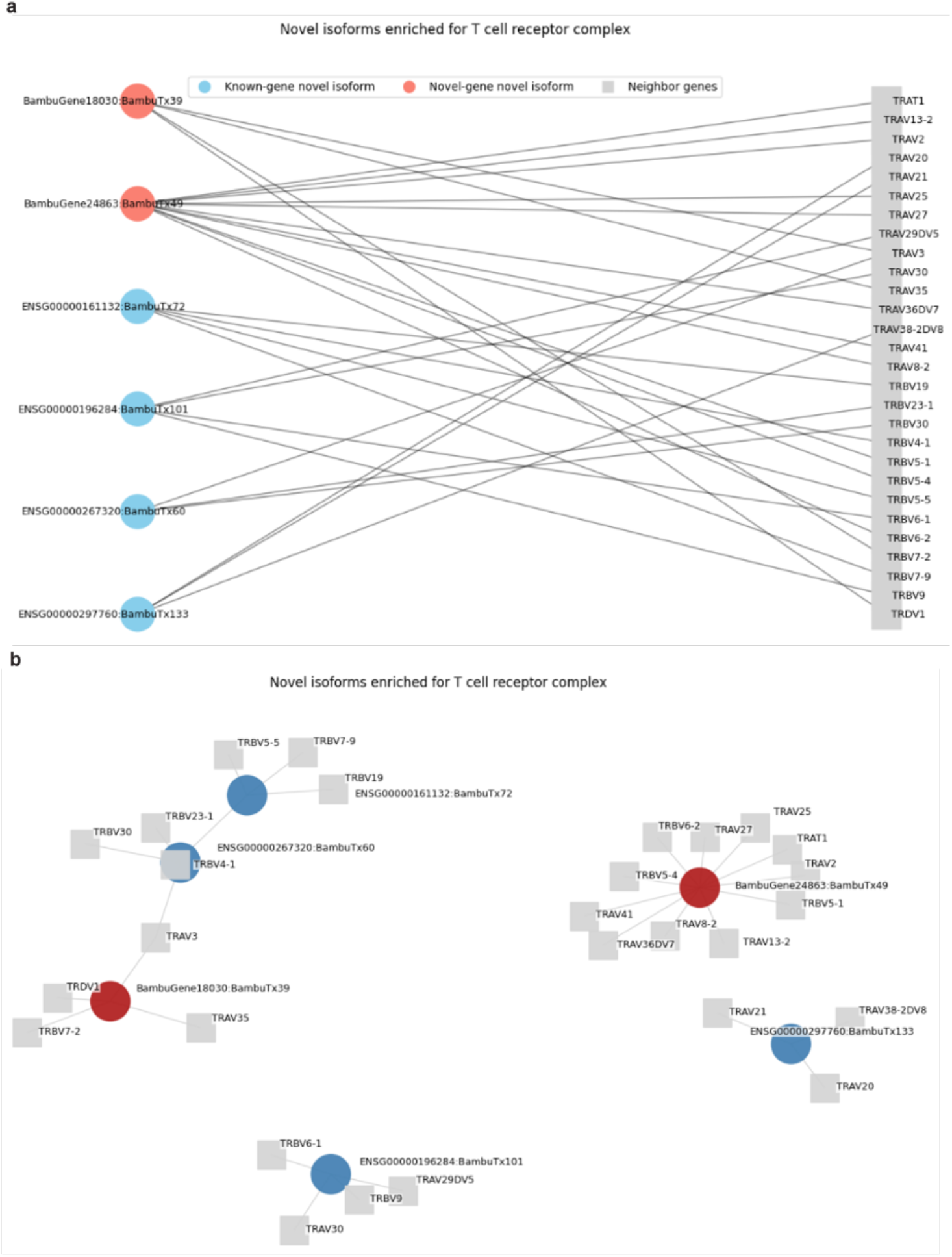
Novel isoforms associated with the T cell receptor signaling complex via their centroid neighborhoods. **(a)** Bipartite network representation of novel isoforms whose centroid neighborhoods were enriched for the GO:CC term, T cell receptor complex (GO:0042101). Nodes represent novel isoforms (left) from previously unannotated (red) and known genes (blue), and centroid neighbor genes localized to the T cell receptor complex (right, grey). Lines connect new isoforms to their neighbor genes based on similarity of cell-type expression profile. (**b)** Network representation of isoform nodes surrounded by immediate associated neighbor genes localized to the T cell receptor complex. Isoform nodes only show overlap of *TRBV* and *TRAV* genes in one instance (*TRAV3*).

*PRODHLP* (*ENSG00000161132*) is categorized as a pseudogene, a gene class recognized as regulating expression at the RNA, DNA, or protein level, in some cases^90–92^. Although pseudogenes remain understudied, some have been implicated in leukocyte development and immune-related disease processes^90^. *SUPT3H* (*ENSG00000196284*) is a nuclear protein-coding gene whose protein functions in RNA polymerase II-mediated transcription regulation as part of multiple transcriptional complexes^93–95^, including the SAGA (Spt-Ada-Gcn5-Acetyltransferase) complex, a regulator of T cell activation^94,96^.

Both genes lacking conventional gene symbols (*ENSG00000267320* and *ENSG00000297760*) are annotated as lncRNAs, a class of genes that participate broadly in health and disease, including tumor suppression^97^ and transcriptional regulation^98^. lncRNAs have been implicated in the regulation of essential immune processes (for review, see ^99^), including immune cell differentiation^100^, activation^101^, and response with cellular lineage- and stimulus-specific expression^102^. Multiple databases have been built to report associations between lncRNAs and their targets, function, and association with disease, which we queried to further contextualize the immune relevance of these genes. Despite neither gene being annotated in general lncRNA databases such as MNDR^103^, EVLncRNAs^104^, and LncSEA^105^, the immune-specific database ImmReg^106^ reported *ENSG00000267320* as being associated with diffuse large B cell lymphoma (DLBC) and thymus cancer (THYM), with a specific association to the TCR signaling pathway, providing external context consistent with the pathway associations identified through our centroid neighborhood analyses. *ENSG00000297760* did not appear in any of these databases.

Because these novel isoforms arise from host genes annotated as potentially regulatory gene types (SAGA complex component, lncRNAs, and a pseudogene), and because their centroid neighbors were enriched for *TRAV* and *TRBV* genes, we hypothesize that these novel isoforms may be associated with regulatory contexts related to *TRAV/TRBV* transcriptional programming, rather than structural components of the TCR complex. This case study demonstrates the specific, hypothesis-generating utility of our centroid neighborhood framework for pathway inference at the level of individual novel isoforms.

## Discussion

Long-read sequencing provides a major advantage to novel isoform detection, rapidly expands the catalog of known isoforms and their biological context in specific diseases and cell types. However, interpreting the relevance of the novel isoforms remains a critical bottleneck. Genomic location alone rarely informs the functionality of novel isoforms, and existing tools for functional interpretation rely on an incomplete gene- and protein-centric view and are therefore insufficient for functional inference. At present, predicting the biological context of novel isoforms is enigmatic before validation and testing via experimental studies. In this study, we present CIPI, a centroid-based framework that addresses this gap by leveraging cell-type expression similarity to generate pathway-informed hypotheses for individual isoforms with or without prior annotation.

In single-cell transcriptomic approaches, centroids are typically used at the cell-level rather than the gene-level, acting as an alternative to KNN for dimensionality reduction^107^, or to assign cell type based on signatures found in existing datasets^108^. In these approaches, distance from the reference centroids is treated as “noise” or batch effects and is discarded or reduced, rather than treated as a signal that could be biologically meaningful^109,110^. Notably, centroid-based approaches have rarely been applied to isoform-level data, where divergence from the canonical isoform may in fact represent true biological variation. CIPI utilizes centroid logic differently, treating variance as a potential biological signal and building a reference gene-set accordingly. Existing annotation resources like DeepGO and the most updated version of the Human Protein atlas, which includes transcript annotations, remain rooted in canonical protein evidence. Using these approaches, novel isoforms remain unannotated, and non-canonical protein-coding isoforms are not independently examined. In contrast, CIPI does not root isoform analyses in their canonical form, instead treating all isoforms as equally relevant to their gene centroid.

The most similar existing approach to CIPI is hdWGCNA, due to their shared use of centroids; CIPI differs in several key ways. In WGCNA-based approaches, the eigengene or eigenisoform summarizes the dominant expression pattern, so isoforms act as participants within a module and any pathway-level context is applied to the module as a whole. This can be a powerful tool to recognizing patterns across several genes and isoforms; however, it does not enable a focused view on individual isoforms. CIPI treats a single isoform as the focal point and builds a neighborhood of genes with similar expression patterns around it. Critically, these centroid neighborhoods are not intended to function as isoform identity classifiers; the objective is not to assign isoforms to dedicated modules. Unlike the modules found in WGCNA, the centroid neighborhood is not a summary of the isoform’s expression—it acts as a reference expression profile assembled from the isoform’s nearest neighbors, using similar expression patterns from genes with known functions to form pathway-informed hypotheses regarding isoform function. In the case of novel isoforms, this centers the novelty and asks which known genes can provide context according to their similarity, allowing for targeted generation of isoform-specific hypotheses.

Since this manuscript focuses on identifying cell-type-specific patterns in both novel and known isoforms and using gene-level pseudo-bulk-by-cell-type analyses to infer functionality, a dataset with sufficient depth was necessary. This framework uses the guilt-by-association principle as the conceptual foundation for the neighborhoods; CIPI assumes that cell-type expression pattern similarity reflects a shared context, which requires accurate cell-type labels and consistent expression differences, rather than technical noise or transient states. For this reason, we chose to use the AutoZI-denoised expression data rather than the raw data. Denoising increases the stability of our cell-type-specific signals, which supports consistent correlation-based neighborhood centroid assembly, but may also reduce the effect of extreme or rare expression patterns. Accordingly, the conclusions formed in this report should be interpreted as proof-of-concept, demonstrating a novel computational strategy to generate hypotheses on the biological enrichment of novel isoforms and understudied genes according to pathway enrichment context, rather than acting as a definitive functional annotation.

Applying CIPI to long-read single-cell data from PBMCs revealed biologically meaningful patterns. Centroids were highly similar to their isoform of interest, exhibiting low residuals for most isoforms and supporting their use as stable reference profiles. Using cell-type marker gene isoforms as a positive control, centroid neighborhoods successfully retained biologically meaningful context, corroborating the cell-type labels from Doyle *et al*.^14^ and supporting the utility of CIPI for pathway-informed contextualization. Intriguingly, although we expected host marker genes to be among the top centroid neighbors of their own isoforms, host genes had a median neighborhood rank of 20, with isoform profiles more closely resembling their neighbor gene centroids than their own host gene. This underscores the limitations of host gene annotation as a proxy for isoform-specific enrichment. Compared to the marker isoforms, we identified less overlap among enriched terms for novel isoforms, with no pathway representing greater than four isoform centroid neighborhoods. This is not unexpected, since the marker isoforms are directly involved in cell-type-specific processes, whereas the host genes for our novel isoforms are more varied in their biological roles. Novel isoforms from known genes converged on biological themes related to mRNA metabolism and translation, whereas novel isoforms from new genes were more varied, exhibiting themes of immune signaling and activation as well as translational regulation, both supported by GO cellular compartment analyses. To illustrate the hypothesis-generating potential of CIPI for individual isoforms, we focused on isoforms enriched for the “T cell receptor” GO:CC term from both known and new genes, including non-coding transcripts, to demonstrate how individual isoforms can be meaningfully contextualized through their neighbor genes in the absence of prior annotation. Together, these results suggest that CIPI can capture biologically relevant signals beyond what is currently accessible through existing annotation-dependent approaches and provide pathway-informed context for hypothesis generation.

As with all pathway enrichment analyses, the context of the tissue is an important consideration for interpreting why certain pathways may be enriched^111–113^. Accordingly, several terms in seemingly unrelated pathways can reflect shared signaling motifs, metabolic shifts, or activation programs that are not exclusive to immune cells but emerge in immune cells under specific conditions. In PBMCs, housekeeping processes such as calcium signaling, developmental cytoskeletal remodeling, and migration may appear as non-immune functions, while being inherently tied to immune cell behavior. A major limitation of using pathway analyses to infer functionality is that enrichment depends on successfully mapping centroid neighbor genes to genes represented in existing annotations, which underrepresents the functional significance of pseudogenes, lncRNAs, and poorly understood loci. For this reason, pathway analyses alone are not sufficient to infer function using the CIPI framework; accurate interpretation requires integration of cellular context, cell-type representation, and relevant biological literature beyond these annotations.

Lastly, these centroid-based similarity and enrichment analyses imply pathway-informed context, but do not directly identify molecular mechanisms. Additional computational evidence, such as protein domain and open reading frame prediction, may provide insight into functional potential based on domains present. Although we demonstrate the utility of CIPI in a PBMC context, the framework is not designed specifically for immune cells and could be applicable in other tissue and cell-type contexts where isoform diversity is high and functional annotation is low, such as the human brain. This framework may also extend to bulk-level datasets with sufficient biological variability across conditions, such as sex, disease status, and tissues type, broadening its applicability beyond single-cell contexts. Applying CIPI across tissue and disease states could provide a systematic means of prioritizing the many isoforms represented in the isonome landscape. Isoforms that demonstrate high cell-type specificity and enrichment convergence across multiple pathway annotation sources represent strong candidates for follow-up investigation.

Hypothesis-generating approaches such as CIPI can act as exploratory analyses, allowing for more targeted follow-up studies for efficient use of research funding. Genetic variation between individuals can influence observed isoform differences, so subsequent studies should be powered to identify conserved isoform splicing patterns across individuals. As long-read single-cell studies expand in size, frameworks such as CIPI provide ways to navigate the isonome and uncover potential isoform-level contributions to health and disease.

## Conclusion

Overall, CIPI provides a thorough way to prioritize newly identified or poorly annotated isoforms, such as lncRNAs and pseudogenes, for validation studies using isoform-specific neighborhood fit, enrichment convergence across databases, and localization prediction. This framework also provides a pipeline to generate hypotheses for molecular functions underlying newly discovered isoforms and under-annotated isoforms. While its inferences depend on the assumption that shared cell-type enrichment patterns reflect a shared biological context, the output is testable hypotheses that can guide targeted validation studies. This approach also offers a practical approach to nominate isoform-level candidates for future mechanistic and translational follow-up, an essential step toward understanding isoform-level contributions to health and disease.

## Acknowledgements

This work has been adapted from a chapter of Patricia Doyle’s doctoral dissertation at the University of Kentucky and is presented here as a preprint for public dissemination.

The authors would like to thank the University of Kentucky Center for Computational Sciences (CCS) and Information Technology Services Research Computing for the use of the Morgan Compute Cluster (MCC) and associated research computing resources.

This work was supported by funding provided by the National Institutes of Health [5T32AG078110-02 to University of Kentucky Sanders-Brown Center on Aging, AG078110 to Doyle; R01AG068331 to Ebbert; GM138626 to Ebbert; NS088555 to Stowe]; Alzheimer’s Association [2019-AARG-644082 to Ebbert]; the BrightFocus Foundation [A2020161S to Ebbert.

## Author Information

### Contributions

P.H.D. conceived and designed the study. M.L.P. implemented the preprocessing pipeline, performed all preprocessing, and carried out initial bulk-level isoform discovery. P.H.D. performed cell filtering, pseudo-bulk- and single-cell-level analyses, and generated all figures. P.H.D. and M.T.W.E. wrote the manuscript.

### Correspondence

Please address any correspondence to Mark T.W. Ebbert.

## Data and code availability

Raw long-read scRNA-seq data utilized for this study are publicly accessible in the NIH Sequence Read Archive (SRA) (accession no. SRR37807425, BioProject no. PRJNA1432415, BioSample no. SAMN56321669).

All code used for analyses is publicly accessible at: https://github.com/UK-SBCoA-EbbertLab/single_cell_centroid_pathway_analyses

## Rigor and Reproducibility

A singularity container was used for most analyses in this study to enable the encasement of specific versions of software in a way that is reproducible. Instructions on accessing the singularity container used for this project can be found at the following GitHub repository: https://github.com/UK-SBCoA-EbbertLab/single_cell_analysis_PBMCs/tree/main.

## Use of generative AI and AI-assisted technologies in the writing process

Author(s) utilized ChatGPT to generate and troubleshoot code for the analyses detailed in this manuscript. All code was thoroughly reviewed for accuracy and validated after working with ChatGPT and the author(s) take full responsibility for the code shared on GitHub.

